# Focal and subtle myelin damage in multiple sclerosis-derived post-mortem human brain slice cultures

**DOI:** 10.64898/2026.05.09.723994

**Authors:** Niels R.C. Meijns, Gema Muñoz González, Sabine Stolker, Bert’t Hart, Bonnie C. Plug, Marianna Bugiani, Öznur Bilir, Kasra Roya-Kouchaki, Wulin Teo, Peter Stys, Sophie Hill, Geert J. Schenk, Gijs Kooij, Benjamin Newland, Antonio Luchicchi

## Abstract

The mechanisms that drive myelin damage as seen in demyelinating disorders such as multiple sclerosis remain incompletely understood. Much of our current knowledge is derived from animal models, but interspecies differences limit their relevance in the context of human pathology and could explain why various promising preclinical therapies failed during clinical translation. Human post-mortem organotypic brain slice cultures provide a unique platform to study human myelin biology, as they preserve genetic, cytoarchitectural, pathological and species-specific context. Here, we evaluated myelin integrity in a human post-mortem brain organotypic slice culture model and experimentally induce focal myelin damage.

Human post-mortem organotypic slices cultures retain key features throughout the culturing period, but exhibit gradual cellular and myelin loss over time. Myelin fibres within the white matter remain detectable and present preserved structural and chemical integrity up to 13 days *in vitro*, indicated by the conserved paranodal and nodal organization and stable myelin spectroscopic signature. Delivery of lysophosphatidylcholine using cryogel scaffolds enables focal drug administration throughout the full depth of the slice with minimal diffusion into surrounding tissue and induces localized demyelination after lysophosphatidylcholine application. Similar focal application of the selective Na_v_1.6 stimulator β-mammal scorpion toxin Cn2 induces subtle myelin destabilization. Overall, our results demonstrate the suitability of a human post-mortem brain organotypic slice culture model as an adequate platform for studying myelin damage in a human disease context.

## Introduction

Myelin sheaths are essential for fast electrical conduction and for the trophic support of enwrapped axons [33]. Myelin sheath disruption is a hallmark of a wide spectrum of neurological conditions, including ageing [9], leukodystrophies [11], Alzheimer’s disease [5], and in particular multiple sclerosis (MS) [15]. Yet, the precise mechanisms that initiate myelin destabilization and drive the transition from subtle structural changes to overt demyelination remain poorly understood.

A subtle yet important feature of early myelin damage is the formation of small focal detachments of the myelin sheath from its enwrapped axon, termed myelin blistering or myelin vacuolization [10, 17]. In animal models of demyelination, myelin blistering precedes frank demyelination [10, 27], suggesting that blister may represent an early sign of instability at the level of axo-myelin interface. In MS, myelin blisters have been observed in MS normal-appearing white matter (WM) and prelesional stages of WM degeneration [17, 18], pointing to their possible role as prodromal indicators of demyelination in the context of this disease. Despite their putative pathogenic significance, the cellular events leading to blister formation and their contribution to disease progression remain largely unexplored.

Much of our current knowledge of myelin biology stems from animal models, which have been invaluable in exploring mechanisms of myelin damage and repair. However, most therapeutic strategies aimed at myelin repair that showed promise in these models ultimately failed in humans [14, 42]. These translational setbacks underscore the limitations of animal systems for capturing the full complexity of human myelin pathology [31] and highlight the need for models that bridge the gap between animal models and clinical studies. Human post-mortem brain organotypic slice cultures (HPMB-OSC) offer a unique opportunity to address this translational gap [16, 28–30, 38, 39]. HPMB-OSCs retain the cellular diversity, cytoarchitecture, and microenvironment of native human brain tissue, while remaining viable for several weeks *ex vivo* [28]. Importantly, HPMB-OSCs retain disease-specific characteristics and allow controlled experimental manipulation, enabling the study of pathological mechanisms and assessment of potential therapies [16, 28, 30]. In a proof-of-concept study, HPMB-OSCs exhibited neuronal activity, expressed virally transduced products, and retained disease-specific pathology features [28]. Bath application of the demyelinating agent lysophosphatidylcholine (LPC) to HPMB-OSCs induced a multicellular response characterized by myelin swelling, macrophage-mediated myelin debris clearance, and reactive astrogliosis [28]. However, myelin integrity in HPMB-OSCs, as well as the extent to which myelin disturbances can be experimentally induced in this model, has not yet been systematically evaluated.

In this study, we first assessed changes in the myelin organization and composition during HPMB-OSCs culture. Subsequently, we instigated focal white matter demyelination by locally applying LPC using cryogel scaffolds, a recently developed strategy previously used in rodent models that allows focal drug delivery [7, 41]. Finally, we induced subtle myelin pathology by creating ionic disbalance using the Na_v_1.6-selective scorpion toxin Cn2. Together, these results establish HPMB-OSCs as a suitable model for the investigation of myelin dynamics, bridging the translational gap between animal studies and clinical research.

## Material and Methods

### Human post-mortem brain specimens

Brain tissue was obtained at autopsy from nine MS brain donors, one Huntington disease patient, one donor with post-traumatic stress syndrome, and one non-demented control (Table 1). For all donors, a tissue block from the superior or middle frontal gyrus devoid of macroscopical abnormalities containing approximately 40% cortex and 60% subcortical WM was rapidly dissected, immersed in ice-cold dissection medium consisting of Hibernate-A (Thermo Fisher, A1247501, US) supplemented with 1% penicillin-streptomycin (Thermo Fisher, 15140122, US) and 2.5 µg/ml Amphotericin-B (Thermo Fisher, 15290018, US), and then promptly transported to the laboratory for further processing. Tissue collection was performed on average with a post-mortem delay of 5.5 hours.

**Table 1.** Donor Characteristics.

| ID | Sex | Age | pH CSF | PMD (h:min) | Diagnosis | Cause of death | Figure |
| --- | --- | --- | --- | --- | --- | --- | --- |
| OS05 | f | 71 | 6,83 | 7:05 | MS | Legal euthanasia | 1, 2 |
| OS06 | m | 54 | 6,67 | 5:45 | MS | Deterioration due to MS | 1, 2 |
| OS07 | m | 53 | 6,52 | 2:50 | Non-MS(HD) | Euthanasia | 1, 2, 3 |
| OS08 | f | 76 | 6,39 | 5:05 | MS | Atrial fibrillation, medication stopped | 1, 2, 3 |
| OS10 | f | 67 | 6,26 | 6:45 | MS | Deterioration due to MS | 1, 2, 4 |
| OS11 | f | 71 | 6,56 | 6:30 | MS | Urosepsis | 4 |
| OS12 | f | 69 | 6,10 | 6:15 | MS | Colon cancer | 4 |
| OS13 | f | 54 | 6,65 | 4:30 | MS | Legal euthanasia | 3, 5 |
| OS14 | f | 73 | 6,42 | 8:10 | MS | Deceased during sleep | 5 |
| OS16 | m | 93 | 6,14 | 5:30 | Non-MS (NNC) | n.a. | 1, 2, 3 |
| OS18 | f | 49 | n.a. | 4:00 | Non-MS (PTSS) | n.a. | 5 |
| OS19 | f | 58 | n.a. | 3:40 | MS | n.a. | 4 |

CSF=cerebrospinal fluid; F=female; HD=Huntington's disease; ID=identifier; M=male; MS=Multiple Sclerosis; NNC=Non-neurological control; PMD=post-mortem delay; PTSS=Post-Traumatic Stress Syndrome, n.a.= not available.

The whole study was approved by the local Medical Ethical Committee of the Vrije University Medical Center (Amsterdam, The Netherlands) and all experiments were performed in accordance with the declaration of Helsinki. For all donors, written informed consent for brain autopsy, the use of the medical data, and permission to use the tissue for research purposes was obtained by the Netherlands Brain Bank (https://www.brainbank.nl/).

### Synthesis of sulfonated cryogels

Cylindrical cryogels were synthesized within 3D printed templates. The templates were designed using Autodesk Inventor software as a cylindrical mold with a depth of 0.5 mm and diameters varying from 2 mm to 4 mm. The molds were printed in PlasCLEAR v2 resin using an Asiga Max 27 printer (Asiga, Australia) with a step size of 50 µm (in the Z-plane). A precursor solution containing polyethylene glycol diacrylate (molecular weight 700, Merck, Germany) and 3-sulfopropyl acrylate (Merck, Germany) at a 1:1 molar ratio was prepared in distilled water at 10% wt/vol, with 2-hydroxy-2-methylpropiophenone used as a photoinitiator at a 1:35 photoinitiator to monomer ratio. The precursor solution was added to the cylindrical molds in the template, which was placed at −20 °C for 30 minutes. A hand-held UV lamp with two 15 W bulbs was then placed in the freezer to irradiate the filled templates for two minutes, polymerizing the monomers to a polymer network. The formed cryogels were removed from the template and washed three times in 10 mL of ethanol, with each wash lasting for at least one hour. The remaining ethanol was then removed and the cryogels were left to dry under vacuum at room temperature and stored dry at room temperature until use.

### Human post-mortem organotypic brain slice cultures

HPMB-OSCs were obtained and cultured based on previously published work.[28] Briefly, collected tissue blocks were mounted on a Vibratome (Leica VT1000S, Germany) containing ice-cold Hibernate-A (Thermo Fisher, A1247501, US) supplemented with 1% penicillin-streptomycin and 2.5 µg/ml Amphotericin B, and 400 μm-thick slices were cut perpendicularly to the cortical surface. Randomly selected slices were immediately fixed in 4% paraformaldehyde (PFA) for 24–48 h and used for the control measurements of day-in-vitro (DIV) 0. The remaining slices were transferred onto semi-permeable membrane inserts (Merck-Millipore, PICM0RG50, Germany) in six-well culture plates containing 1 ml of normal slice culture medium, which consisted of 50% Minimum Essential Medium (Thermo Fisher, 32360026, US), 25% Earle’s Balanced Salt Solution (Thermo Fisher, 24010043, US), 25% heat-inactivated horse serum (Thermo Fisher, 26050088, US) supplemented with 1% GlutaMAX-I (Thermo Fisher, 35050038, US), 1% penicillin-streptomycin, 1.25 µg/ml Amphotericin B, and 2.6 mg/ml D-Glucose (45% in dH2O; Sigma-Aldrich, G8769, Germany).

At DIV 0 (i.e. before culturing conditions), 5, 9, 13 and 20, slices were fixed with 4% PFA for 24–48 hours and stored until further processing on 0.05% sodium azide in PBS at 4 ºC. Slices were either embedded in paraffin or frozen depending on the research question. Slices used for assessing cyto- and myeloarchitecture during the culturing period were paraffin-embedded and sliced into 5 μm-thick sections using a microtome (Leica Biosystems, Germany). Slices used for Nile Red spectroscopy and for assessing drug delivery effects were cryoprotected overnight in 10% (w/v) sucrose (Sigma-Aldrich, 1604, Germany) solved in PBS at 4 ºC. Each slice was then placed over a 3×3 cm steel platform and enclosed within a modified Tissue-Tek® Cryomold (Sakura Finetek, 62534-25, 25×20×5 mm, US) from which the base had been removed, leaving only the four lateral walls as a frame. The steel plate was then placed over crushed dry ice and Optimal Cutting Temperature (Cell Path, 361603, UK) was rapidly added, allowing to quickly and uniformly freeze and embed the tissue in a flat orientation while preserving the culture plane. Slices were then sectioned parallel to the culture plane using a cryostat (Leica Biosystems, Germany) into 25 or 20 µm-thick sections (for Nile Red spectroscopy and drug delivery assessment, respectively) and mounted on Superfrost™ Plus slides (Menzel, 6319483, Germany).

#### LPC administration

Cryogels were sterilized with 95% ethanol and desiccated prior to use by drying at room temperature inside a sterile biosafety cabinet. Subsequently, cryogels were loaded with either egg yolk-derived LPC (10 or 15 mg/ml in PBS, Sigma-Aldrich, L4129, Germany) or sterile PBS. At DIV 7, LPC- and PBS-loaded cryogels were carefully placed onto the subcortical WM region of the brain slices at positions equidistant from the adjacent grey matter (GM), using sterile forceps, ensuring maximum distance between individual gels. After 18 h, designated as day post-lesion (DPL) 0, all gels were carefully removed using forceps, and the culture medium was refreshed. At DPL 3, slices were fixed, cryoprotected, frozen, embedded, and sectioned as indicated above.

For fluorescent LPC delivery, 3 mm-diameter cryogels were loaded with a freshly prepared mixture consisting of 10 mg/mL LPC solution in PBS, in which 15% (w/w) of the LPC was BODIPY-labeled (Top Fluor® LPC, Avanti Polar Lipids, US). Gels were loaded with 10 µL, corresponding to the empirically determined minimum reconstitution volume. For bath LPC application, freshly prepared culture medium containing 1 mg/mL LPC with 15% BODIPY-labeled LPC was used. After 18 h, gels were removed if applicable, and medium was refreshed. Slices were fixed either immediately (i.e. at DPL 0) or at DPL 3, subsequently cryoprotected, frozen and embedded, and cut perpendicularly to the vibratome slicing plane into 20 µm-thick slices.

#### Cn2 scorpion toxin administration

Using the same procedure described above, synthetic, single chain β-scorpion neurotoxic peptide (STC-060, Alomone Labs, US) was dissolved in ddH20 and loaded into desiccated 2 mm-diameter cryogels to a final concentration of 140 nM [32] using the empirically determined minimum reconstitution volume (2.5 μl). At DIV 7, PBS- and Cn2-loaded cryogels were applied over the WM of HPMB-OSCs for 12 h, and slices were fixed in 4% PFA for 8 h after removal of cryogels.

### Nile Red spectroscopy

Nile Red (9-(diethylamino)-5H-benzo[a]phenoxazin-5-one, Sigma Aldrich, 72485, Germany) was prepared at 5 mM concentration in DMSO and stored at 4 °C. To assess the nature of Nile Red solvatochromism, Nile Red was dissolved in several solvents with varying dielectric polarities in glass capillaries and excited with 445 nm. Emission spectra were recorded following excitation with a 445 nm laser.

For biological specimens, Nile Red was diluted to a final concentration of 30 µM in PBS. Fixed-frozen 25 µm tissue sections were stained with Nile Red for 10 min and subsequently rinsed for 5 min in PBS. The stained tissue sections were then submerged in PBS solution on a glass microscope slide and sealed with silicon grease to enable imaging on an inverted microscope. The procedure for Nile Red spectroscopy was carried out based on a previously published protocol [34], with some adaptations. All images were collected following excitation with a 445 nm laser. Spectral fluorescence images were recorded using an inverted Nikon A1R spectral confocal microscope equipped with a Plan Apo VC 20× objective (0.75 NA, Nikon, Japan). Spectrometer resolution was 10 nm with 26 channels ranging between 490 nm and 750 nm. All spectra were interpolated to 400 data points using cubic splines, and each spectrum was then normalized for comparison and spectral decomposition using ImageTrak image analysis software (http://stysneurolab.org/imagetrak/). For visualization of multichannel spectral images, “true color” renditions were rendered to approximate what the naked eye would see if looking at the samples during microscopy. For spectral decomposition, two bracketing basis spectra were used to define the minimum and maximum polarity values represented by the most blue- and most red-shifted spectra, respectively. To generate pseudocolor images, a nonlinear transformation algorithm assigned a numerical index between 0 and 1 depending on the fit of a given pixel to the pair of bracketing reference spectra. This pixel was then assigned a pseudocolor ranging from violet (index 0) to red (index 1); nonpolar features in the image are therefore represented by cooler colours, while more polar elements appear in hotter colours. The polarity index was defined as this numerical index produced by the nonlinear fitting. To select myelin for data analysis, images were acquired at 5× to 10× zoom and myelin was based on intensity after manually segmenting out autofluorescent structures in order to minimize influence of autofluorescence on spectral analysis.

### Histology and immunohistochemistry

#### Luxol Fast Blue (LFB)

Paraffinized sections were first placed on a heating plate for 45 minutes at 57 ºC, incubated with 3×100% xylene substitute for 10 minutes and 2×100% ethanol for 5 minutes per step. Paraffin-embedded and frozen sections were incubated with filtered 0.1% LFB (Gurr, Electron Microscopy Sciences, Hatfield, PA, USA) solved in 96% ethanol overnight at 56 ºC. Subsequently, sections were briefly rinsed in 96% ethanol and ultrapure water and differentiated in 0.05% lithium carbonate and 70% ethanol until the WM and GM could be macroscopically distinguished. Finally, sections were rinsed with ultrapure water, dehydrated in an ethanol (2×96% ethanol, 2×100% ethanol, 2 minutes per step), and xylene series (3×100% xylene, 2 minutes per step) and mounted with Entellan (Merck, Germany).

#### Immunohistochemistry

For immunohistochemistry, slices were deparaffinized by placing them on a heating plate at 57ºC for 45 minutes and incubating them for 5 minutes each with 3×100% xylene substitute, 2×100% ethanol, and 90% to 80% and 70% ethanol. Antigen retrieval was conducted for 30 minutes in Tris/EDTA buffer pH = 9 in a steam cooker, after which the slices were cooled to room temperature for approximately 40 minutes. Fixed-frozen slides underwent an antigen retrieval phase by placing them in Tris-EDTA for 15 minutes in a steam cooker and subsequently cooled down to room temperature for 15 minutes. The following steps were conducted in a similar fashion for both paraffin-embedded and frozen sections. Sections were incubated with 3% normal donkey serum (Jackson, 017-000-121, US) in TBS containing 0.5% Triton X-100 (TBS-T, Merck, 8603, Germany) for 30 minutes In the case of the Na_v_1.6/CASPR-staining, incubations with primary anti-Caspr1 antibody was preceded by endogenous peroxidase blocking using 1% H_2_O_2_ (Merck, 1.07209.1000, Germany) for 30 minutes. All sections were incubated overnight at 4 °C with the corresponding primary antibodies (**Table 2**), which were all diluted in 3% normal donkey serum in 0.5% TBS-T. Then, sections were washed 3×5 minutes with TBS, and subsequently incubated with appropriate Alexa Fluor®-conjugated secondary antibodies (**Table 2**), washed 3×5 minutes in TBS in the dark, counterstained with DAPI (Sigma, D9542, Germany) 1:1000 in TBS for 5 minutes, briefly washed with TBS, and mounted with Mowiol-Dabco (Sigma, 81381 and 290734, Germany). In the case of slides stained with anti-NeuN, autofluorescence was quenched by incubating the slides with Sudan Black B (0.1% in 70% ethanol) for 10 min, followed by a 2×5 min wash with ultrapure water.

**Table 2.**
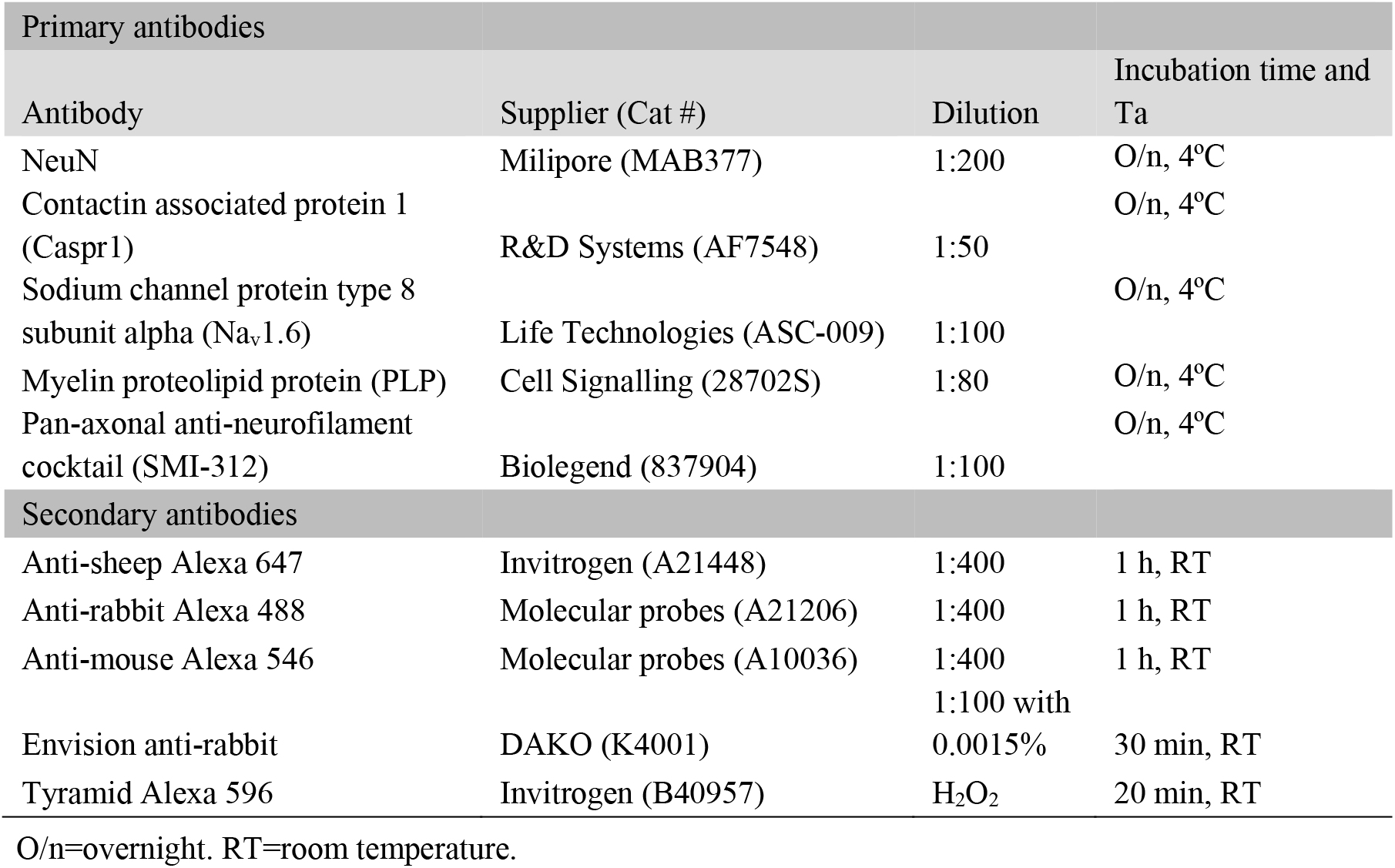
List of antibodies, dilutions, and suppliers.

#### Image acquisition

For bright-field image analysis, images were acquired with whole-slide scanner (Vectra Polaris, 20× objective, Akoya Bioscience, US). For assessing cellular density, NeuN-stained sections were imaged with an OLYMPUS VS200 slide scanner with a numerical aperture of 0.4 using a 10× objective (DAPI filter: Ex = 378 nm, Em = 432 nm, exposure = 10 ms; FITC: Ex 474 nm, Em 515 nm, exposure = 1.00 s; Cy3 for AF; Ex 554, Em 595 nm, exposure = 500 ms). Nodes of Ranvier, and axonal and myelin signal after LPC and Cn2 treatment were imaged using a Leica TCS SP8 (Leica Microsystems, Germany) and an HC PL APO CS2 40× oil objective lens. Fluorophores were excited at their respective optimal wavelengths using a tuneable pulsed white laser and signals were detected by gated hybrid detection using standard mode with pixel sizes 101 nm (nodes of Ranvier) or 142.05 nm (myelin). HPMB-OSCs WM was imaged in tiles of ∼1050 μm^2^ per treatment condition per slide (myelin) or in Z-stacks of 4 μm (step size 0.5 μm) at ∼1000 μm^2^ per slide (nodes of Ranvier). Images were background subtracted and when applicable maximum intensity projections were created using ImageJ Fiji (National Institute of Health, US).

#### Image analyses

##### Nodes of Ranvier

Nodal density was calculated using Na_v_1.6-positive puncta as a proxy for nodes in a subset of cases with comparable background fluorescence. Using max-projected micrographs, nodes bilaterally flanked by Caspr1-positive paranodes were traced with the segmented line function in ImageJ and spline-interpolated and re-mapped to virtually straighten the axon on a pixel-intensity-based fashion.[12] A total of 13–165 axonal segments encompassing the nodal and paranodal regions we traced per HPMB-OSC, with n=3–6 cases per DIV. Nine consecutive, horizonal intensity profiles of Na_v_1.6 and CASPR1 signal were generated and averaged per axonal segment. The start and end points of node and paranodes were determined using the half-maximum values of Na_v_1.6 and Caspr1 signal, respectively, and were subsequently used to calculate nodal length and the extent of Caspr1 intrusion into the node.

##### LPC

Axonal segments running in parallel to the sectioning plane and without signs of ongoing overt degeneration were traced and straightened as described above. Axons were selected from areas underneath the PBS- and LPC-loaded cryogels area in a blinded manner. Consecutive line scans perpendicular to the axonal axis were made to assess the myelination profile, using a custom-made script (available upon request). For each condition (PBS or LPC), approximately 100 axons were traced from three independent cases, yielding total traced lengths of 882.1 µm and 951.1 µm for the LPC and PBS conditions, respectively.

##### Myelin after Cn2

Similar to the analysis of the nodes of Ranvier, myelin blistering after Cn2 treatment was assessed by tracing and isolating axons using ImageJ’s ‘Segmented line’ and ‘Straighten’ functions to create separate TIFFs for each image. All axons per case and condition were then aligned in a composite image for visual scoring of myelin swelling and blistering by three individuals. The swelling width was measured using a line scan perpendicular to the axonal axis at the point of maximum swelling diameter.

### Statistical analysis

For comparisons of multiple time points, a one-way analysis of variance (ANOVA) was performed, followed by Dunnett’s post hoc test to compare each time point against baseline (DIV 0). For pairwise comparisons between treatment conditions, a paired two-tailed t-test was used. Statistical significance was defined as p < 0.05. All analyses were conducted using GraphPad Prism 10.3.0 (GraphPad Software, Boston, US).

## Results

### Cellular density decreases in HPMB-OSCs are regionally distinct

We first assessed whether HPMB-OSCs retain the principal cytoarchitectural features of viable cortical tissue over time. Rapidly excised cortical blocks were sectioned into 400-μm slices and maintained on semi-permeable inserts for up to 20 DIV (**Fig. 1a**). Tissue was obtained from donors without demyelination or overt inflammatory pathology (Table 1). Cytoarchitecture and neuronal density were evaluated in sectioned cultures immunolabelled for NeuN and counterstained with DAPI (**Fig. 1b**). In the grey matter (GM), total cellular density progressively decreased during the culture period (**Fig. 1d**). Cortical layering remained distinguishable at early timepoints (e.g., DIV 5; **Fig. 1c**), but cell densities fell significantly at later stages, indicating progressive cellular loss. The proportion of NeuN-positive neurons relative to total nuclei also changed over time, with a marked increase at DIV 20 compared to baseline (**Fig. 1e**), suggesting that neurons are relatively more preserved than non-neuronal cells during prolonged culture. WM exhibited a similar overall decline in total cellular density, although the onset of this reduction was delayed compared with GM (**Fig. 1d**). Significant decreases were observed only at later timepoints, consistent with region-specific temporal dynamics in vulnerability. Together, these findings demonstrate that HPMB-OSCs undergo a time-dependent reduction in overall cell density, with GM and WM showing distinct trajectories of cellular change during *ex-vivo* maintenance.

**Fig. 1.**
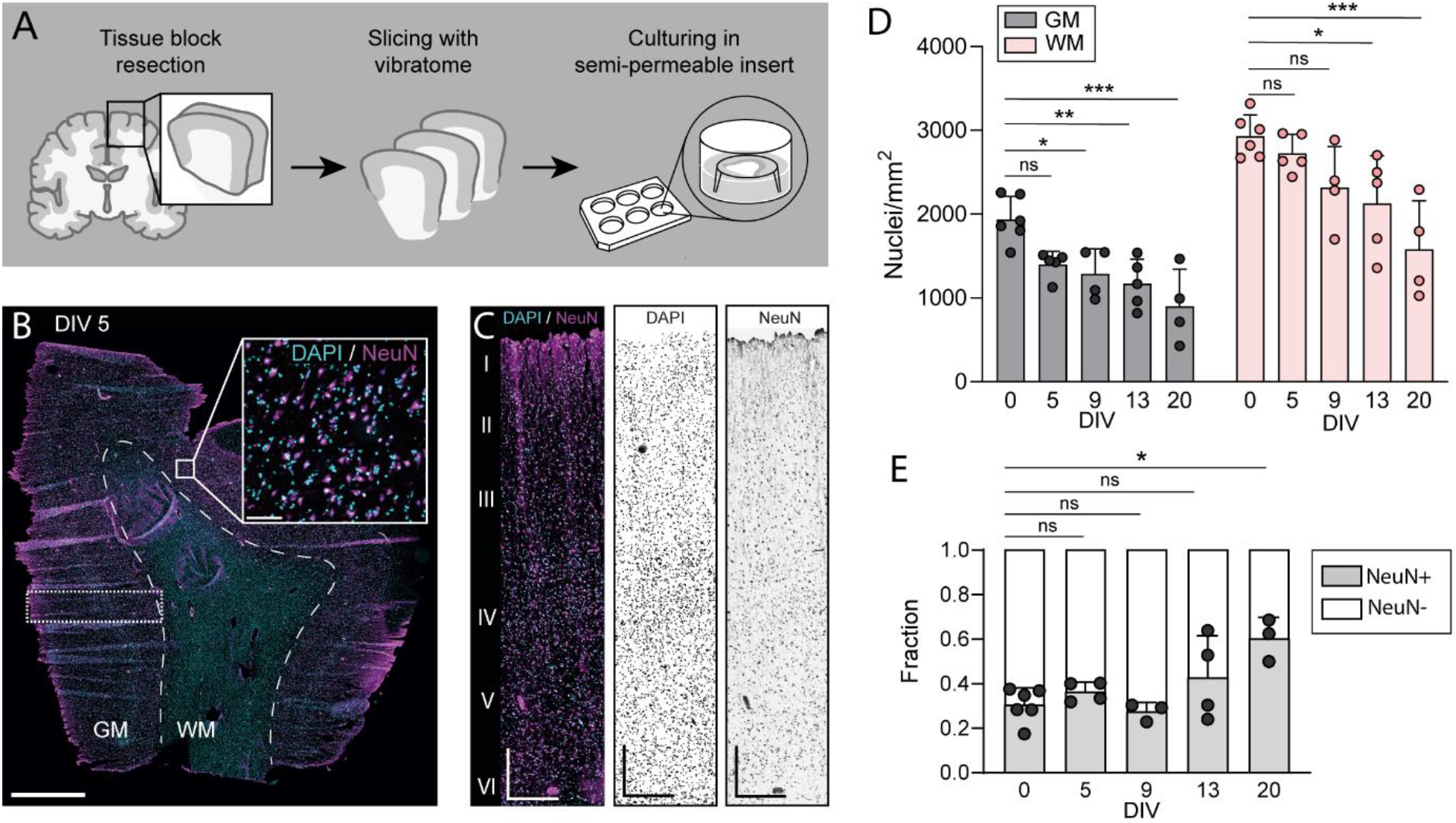
Cellular density in HPMB-OSCs in grey and white matter differentially decrease slowly over time. a. Schematic overview of HPMB-OSC preparation. A 1.5 × 1.5 cm cortical tissue block (superior or middle frontal gyrus) was dissected at autopsy, sectioned at 400 μm, and cultured on semi-permeable inserts. b. Confocal micrograph of a whole HPMB-OSC at DIV 5. Scale bar: 2.5 mm (overview), 500 μm (inset). c. Confocal image of cortical grey matter (GM; region indicated in B) showing preserved cortical layering at DIV 5. Scale bar: 500 μm. d. Cellular density dynamics in GM (left) and WM (right) during the culturing period (N = 4–6 slices per timepoint). GM exhibited a significant overall reduction in cellular density across time (F(4, 19) = 8.69, p < 0.001 (one-way ANOVA)). Post hoc Dunnett’s test showed decreases relative to DIV 0 at: DIV 5: p = 0.505 (ns), DIV 9: p = 0.128, DIV 13: p = 0.021, DIV 20: p < 0.001. WM also showed a significant overall reduction (F(4, 19) = 7.13, p = 0.0011 (one-way ANOVA)). Post hoc Dunnett’s test indicated declines at: DIV 13: p = 0.02, DIV 20: p = 0.004. Earlier timepoints n.s. e. Fraction of NeuN-positive cells relative to total DAPI-positive nuclei during culturing (N = 3-6). NeuN fraction changed significantly over time (F(4, 15) = 5.52, p = 0.0076 (one-way ANOVA)). Post hoc Dunnett’s test revealed a significant increase at: DIV 20 vs. DIV 0: p = 0.0038.

### Myelin fibres in HPMB-OSCs retain healthy structural properties throughout culture

Next, we assessed macroscopic myelin integrity throughout the culturing period using PLP immunohistochemistry and LFB staining. Myelin architecture in the cortex of HPMB-OSCs was largely preserved at DIV 5, although myelin swellings were abundant in the WM, which may reflect intrinsic tissue pathology rather than culture-induced changes (**Fig. 2a**). Individual myelinated fibres projecting into the WM remained visible throughout the entire culture period (**Fig. 2b, c**). However, the boundary between GM and WM gradually became less distinct over time (**Fig. 2b, c**), suggesting partial loss or disorganization of WM myelin.

**Fig. 2.**
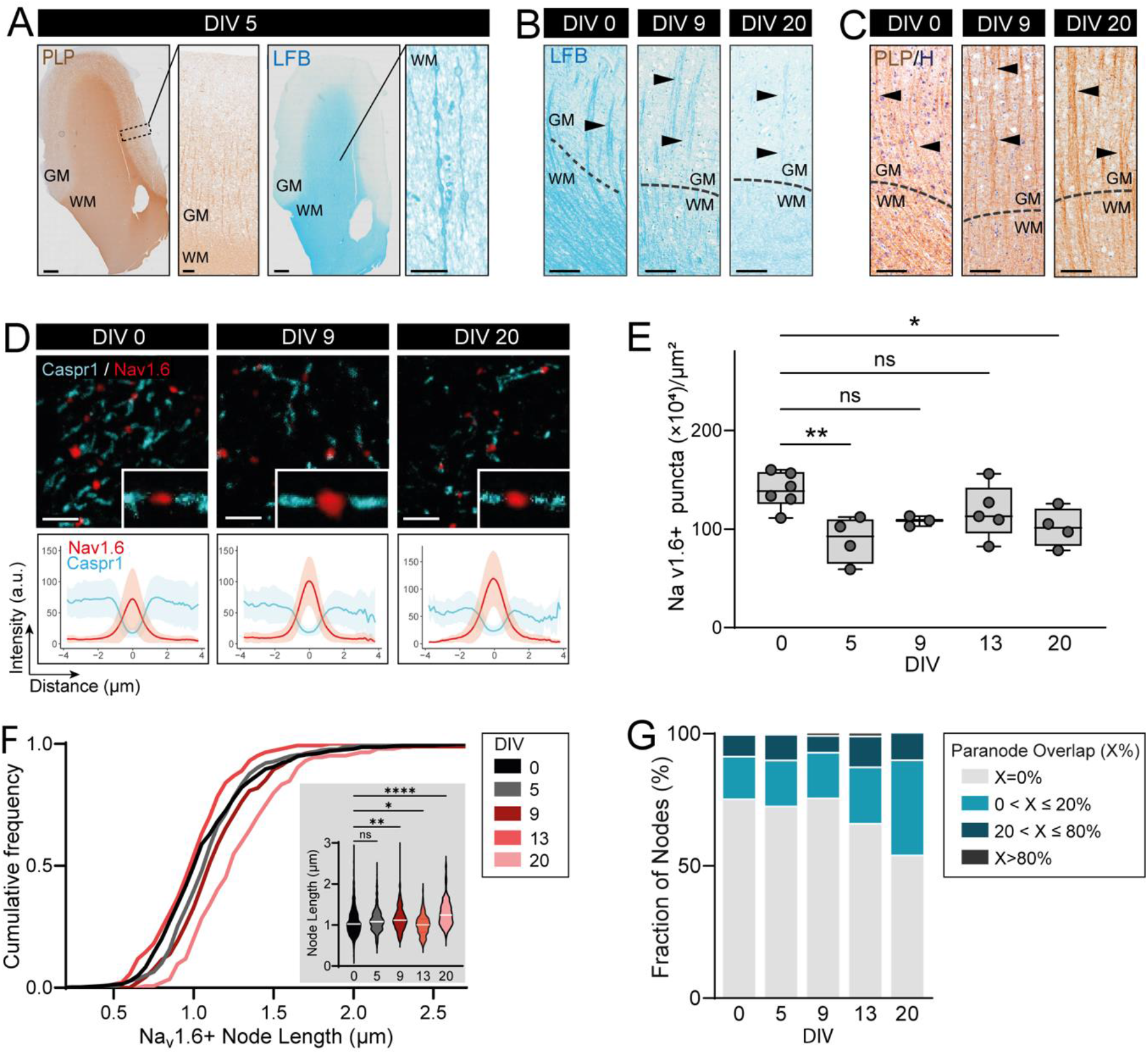
Relative preservation of myelin structure in HPMB-OSCs throughout the culturing period. a. Micrograph showing preserved macroscopic myelin integrity in cortex at DIV 5. Scale bar: 1 mm (overview), 100 μm (inset). b. High-magnification LFB-stained HPMB-OSC showing myelinated fibres (arrowheads) extending from GM into WM up to DIV 20. Scale bar: 100 μm. c. High-magnification PLP-immunolabelled section showing GM-to-WM myelinated fibres (arrowheads) visible throughout culture. d. Representative images of nodes of Ranvier (Na_v_1.6, red) and paranodes (CASPR1, cyan), with averaged intensity profiles across all traced segments. Scale bar: 5 μm. e. Na_v_1.6-positive nodal puncta per 10,000 μm^2^ across DIV 0–20 (individual points represent biological replicates available at timepoints). One-way ANOVA: F(4,17) = 4.073, p = 0.017; Dunnett’s post hoc vs DIV 0: DIV 5, p = 0.0062; DIV 20, p = 0.0429 (other comparisons not significant). f. Cumulative distribution and mean node length across DIV 0–20 (n = 3–6 cases per DIV, 13–165 nodes per case). One-way ANOVA: F(4,1471) = 19.71, p < 0.0001; Dunnett’s post hoc vs DIV 0: significant increases from DIV 9 onward (p < 0.0001). g. Fraction of nodes exhibiting 0%, slight (0–20%), moderate (20–80%), or complete (>80%) paranodal overlap (defined as Caspr1+/Na_v_1.6+ double-positive length relative to total Na_v_1.6-positive nodal length). Fisher’s exact test: DIV 0 vs DIV 5, 9, and 13 not significant; DIV 0 vs DIV 20, p < 0.001.

To further investigate myelin health, we examined the presence of intact paranodal and nodal structures across the culturing period. Na_v_1.6-immunoreactive nodes flanked by CASPR1-positive paranodes were observed from DIV 0 up to DIV 20 (**Fig. 2d**). The density of Na_v_1.6-positive nodal puncta decreased over time (**Fig. 2e**), while node length progressively increased, becoming most prominent at DIV 20 (**Fig. 2f**). This elongation, consistent with impaired axo-myelinic attachment at the paranodes, aligns with features of nodal disruption described in demyelinating conditions [2–4, 26]. We next quantified the degree of paranodal–nodal overlap as an additional readout of structural integrity. At DIV 0, the majority of nodes displayed no overlap between Na_v_1.6-positive nodal segments and Caspr1-positive paranodes, with only a small fraction exhibiting slight or moderate overlap (**Fig. 2g**). This distribution remained stable up to DIV 13. By DIV 20, however, a shift toward increased overlap was observed, indicating displacement of nodal and paranodal proteins and suggesting progressive weakening of paranodal junctions with extended culture.

### Spectral analysis rapport stable WM tissue composition in HPMB-OSCs

To further assess the stability of HPMB-OSCs throughout culture, we employed Nile Red spectroscopy as a proxy for overall tissue composition. Nile Red is a lipophilic fluorescent dye whose emission spectrum shifts depending on the polarity of its microenvironment (**Fig. 3a**), allowing detection of compositional changes in biological tissues [34, 35]. Because GM and WM differ substantially in their lipid, protein, and water content -with WM being enriched in lipid-dense myelin and GM containing more water- and protein-rich cellular components-their spectral profiles are characteristically distinct, with GM exhibiting a more red-shifted emission (**Fig. 3b**). This distinction was maintained up to DIV 13, as visualized through pseudocolor polarity maps (**Fig. 3b**), indicating preservation of broad tissue composition. WM displayed a progressive red shift over time, which may reflect subtle myelin loss and expansion of extracellular water-rich spaces.

**Fig. 3.**
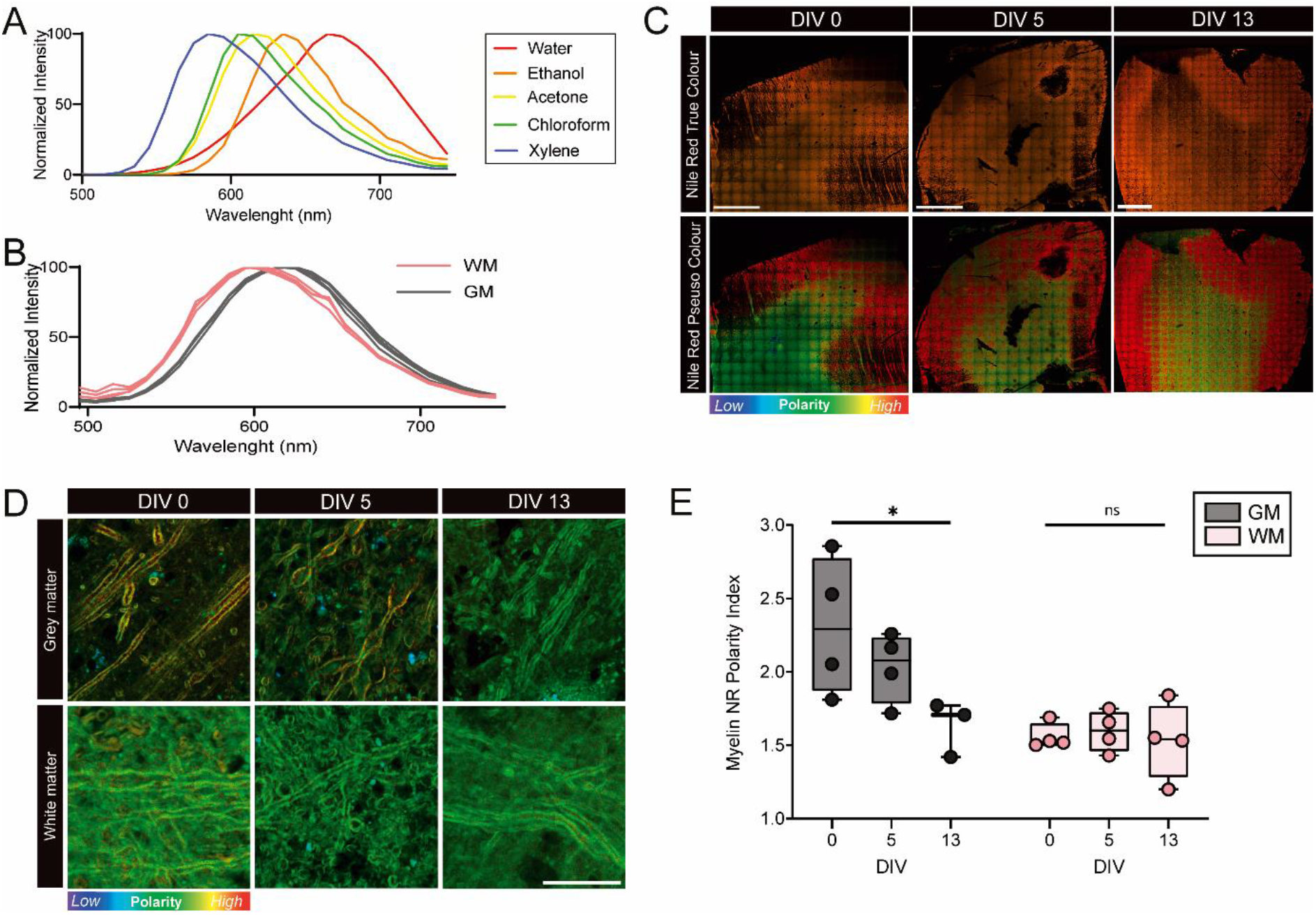
Quantitative spectral analysis of HPMB-OSCs tissue composition with Nile Red solvatochromism. a. Normalized Nile Red fluorescence emission spectra for solvents with different dielectric constants. b. Normalized Nile Red fluorescence emission spectra for WM and GM at DIV 0 from four individual cases. c. Micrographs of whole cryosectioned HPMB-OSCs stained with Nile Red, displayed in true colour (top) and pseudocolor (bottom) at DIV 0, 5, and 13. Pseudocolor contrast reflects polarity of the Nile Red microenvironment, with warmer colours indicating more polar environments and cooler colours indicating more apolar environments. Scale bar: 2500 μm. d. High-magnification pseudocolor images of myelinated axons in WM and GM regions at DIV 0, 5, and 13. Scale bar: 15 μm. e. Mean polarity index derived from Nile Red spectra of segmented myelin in GM and WM. Individual datapoints represent cultured slices from different cases. Comparisons between GM and WM polarity at DIV 0: unpaired t-test with Welch’s correction, WM = 1.56 ± 0.09, GM =2.31 ± 0.47; t(3.2) = 3.15, p = 0.047. Changes across time: GM, Kruskal–Wallis H = 5.60, p = 0.048; WM, Kruskal–Wallis H = 0.500, p = 0.815.

Given the high lipophilicity of Nile Red and its preferential localization to lipid-rich environments, we next assessed myelin compartment polarity using intensity-based segmentation of myelinated fibres. At DIV 0, GM myelin showed higher and more heterogeneous polarity than WM myelin, consistent with previous observations (**Fig. 3e**). During culture, GM myelin polarity decreased, whereas WM polarity remained stable, suggesting overall preservation of WM myelin chemical properties in HPMB-OSCs during the culture period.

### Demyelination by focal administration of LPC in HPMB-OSC white matter

Previously, bath delivery of LPC to HPMB-OSCs was shown to elicit myelin swelling and engagement of microglia/macrophages in myelin phagocytosis, although demyelination was not reported [28]. Bath application of LPC or other compounds introduces two limitations: (1) the entire slice is exposed, and responses to the drug can differ depending on the precise location within the slice (e.g., subcortical vs. deeper WM), and (2) individual slice cultures are required for treated and control conditions, increasing tissue usage and inter-culture variability. In contrast, cryogel-based focal delivery enables sharply localized insults surrounded by NAWM, more closely mimicking focal MS pathology than bath application.

Therefore, we used cryogels to focally deliver LPC or vehicle (PBS) focally to the same HPMB-OSC slice (**Fig. 4a**). To evaluate LPC penetration, cryogels were loaded with LPC and fluorescently labelled LPC (15% w/w) and placed onto WM regions for 18 h. Bath application resulted in diffuse but uneven LPC distribution immediately after treatment, with substantially reduced signal at DPL 3 (**Fig. 4b**, top). In contrast, cryogel-mediated delivery produced sharply confined lateral spread and deep axial penetration through the slice thickness, with minimal lateral diffusion at both DPL 0 and DPL 3 (**Fig. 4b**, bottom). LFB-stained cryosections showed that PBS-loaded cryogels did not induce detectable macroscopic demyelination at DPL 3 (**Fig. 4c**), whereas LPC-loaded cryogels produced marked, spatially restricted demyelination beneath the gel. Confocal imaging confirmed this, revealing intact axons but substantial PLP loss directly under LPC-loaded gels at DPL 3, while adjacent areas and PBS-treated regions maintained PLP signal (**Fig. 4d**). Tracing of morphologically intact axons and corresponding line scans demonstrated clear myelin signal surrounding axons in PBS-treated areas, whereas PLP signal was absent in axons beneath LPC-loaded gels (**Fig. 4e**). Quantification of PLP intensity further confirmed a reduction in myelin signal in LPC-treated regions, supporting successful induction of focal WM demyelination via topical cryogel-mediated LPC delivery.

**Figure 4:**
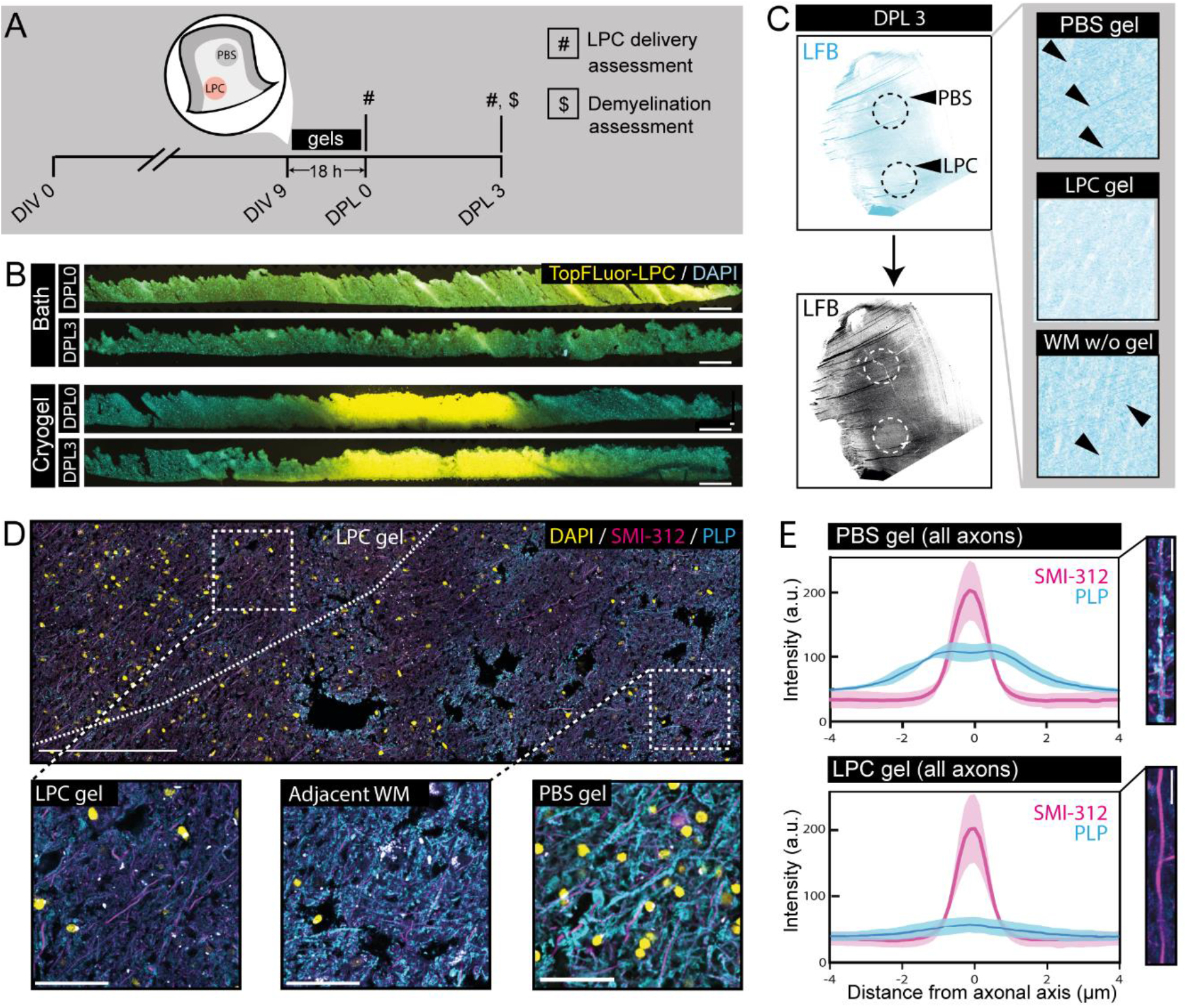
Focal LPC delivery using cryogels induces focal white matter demyelination in HPMB-OSCs. a. Schematic overview of the experimental procedure. Cryogels loaded with LPC or PBS were placed onto WM of HPMB-OSCs for 18 h, with LPC penetration and demyelination assessed at DPL 0 and DPL 3. b. Confocal micrographs of HPMB-OSCs after LPC delivery via bath application (top; 1 mg/ml LPC, 15% TopFluor LPC) or a 3-mm cryogel (bottom; 10 mg/ml LPC, 15% TopFluor LPC) at DPL 0 and DPL 3. Scale bar: 400 μm. c. Luxol Fast Blue staining at DPL 3 showing preserved staining under PBS-loaded cryogels and reduced staining under LPC-loaded cryogels (10 mg/ml), consistent with localized demyelination. Original and grayscale images shown. d. Confocal micrographs of HPMB-OSCs treated with a 3-mm cryogel loaded with LPC (15 mg/ml). DAPI (yellow), SMI-312 (magenta), PLP (cyan). Dashed outline indicates gel position. Scale bar: 250 μm and 50 μm (magnifications). e. Average linescans of straightened axons from HPMB-OSCs (N = 3) at DPL 3 following treatment with LPC-loaded and PBS-loaded cryogels. Approximately 100 axons were traced per condition, spanning 882.1 μm (LPC) and 951.1 μm (PBS). Mean maximum PLP intensity (±2 μm around axon heartline): LPC, 51.6 ± 11.2; PBS, 103.4 ± 8.5; unpaired t-test, p = 0.04. Scale bar: 10 μm.

### Cryogel application of scorpion toxin Cn2 causes alters myelin integrit

After confirming effective focal LPC-induced demyelination and previously observing neuronal activity at DIV 11 in HPMB-OSCs [28], we asked whether sodium channel activity could contribute to myelin damage and myelin blister formation in HPMB-OSCs. We previously observed myelin swelling in early stages of myelin damage and found that manipulation of sodium channels was able to dictate myelin swelling fate [1, 21]. Because oligodendrocytes buffer ions to maintain homeostasis at the axon– myelin interface [19, 22–24], we sought to induce myelin swelling by increasing Na^+^ flux. To this end, we applied the β-mammal neurotoxic peptide Cn2, which induces persistent sodium influx through Na_v_1.6 channels at the nodes of Ranvier [32].

At DIV 7, HPMB-OSCs were treated with Cn2- and PBS-loaded cryogels for 12 h, and myelin and axonal integrity were examined eight hours later by confocal imaging of PLP- and SMI-312–stained slices (**Fig. 5a, b**). Non-cultured slices (DIV 0) were also stained to assess baseline axo-myelinic structure. Myelinated axons lacking overt degeneration were traced in PBS- and Cn2-treated regions and were scored for myelin blisters (local detachments of myelin from the axon) and blebs (combined axonal and myelin swelling; **Fig. 5c**). Baseline blister and bleb frequencies varied across donors and did not increase during culture (**Fig. 5d, e**). Likewise, no differences were observed between PBS- and Cn2-treated regions in the number of blisters or blebs per axonal millimetre, indicating that the selected dose and exposure duration of Cn2 did not induce new swellings. Interestingly however, blister width was greater in Cn2-treated regions than in PBS controls (**Fig. 5f**), suggesting that perturbation of sodium homeostasis may in this case subtly alter myelin structural integrity without increasing the number of axo-myelinic detachments.

**Fig. 5.**
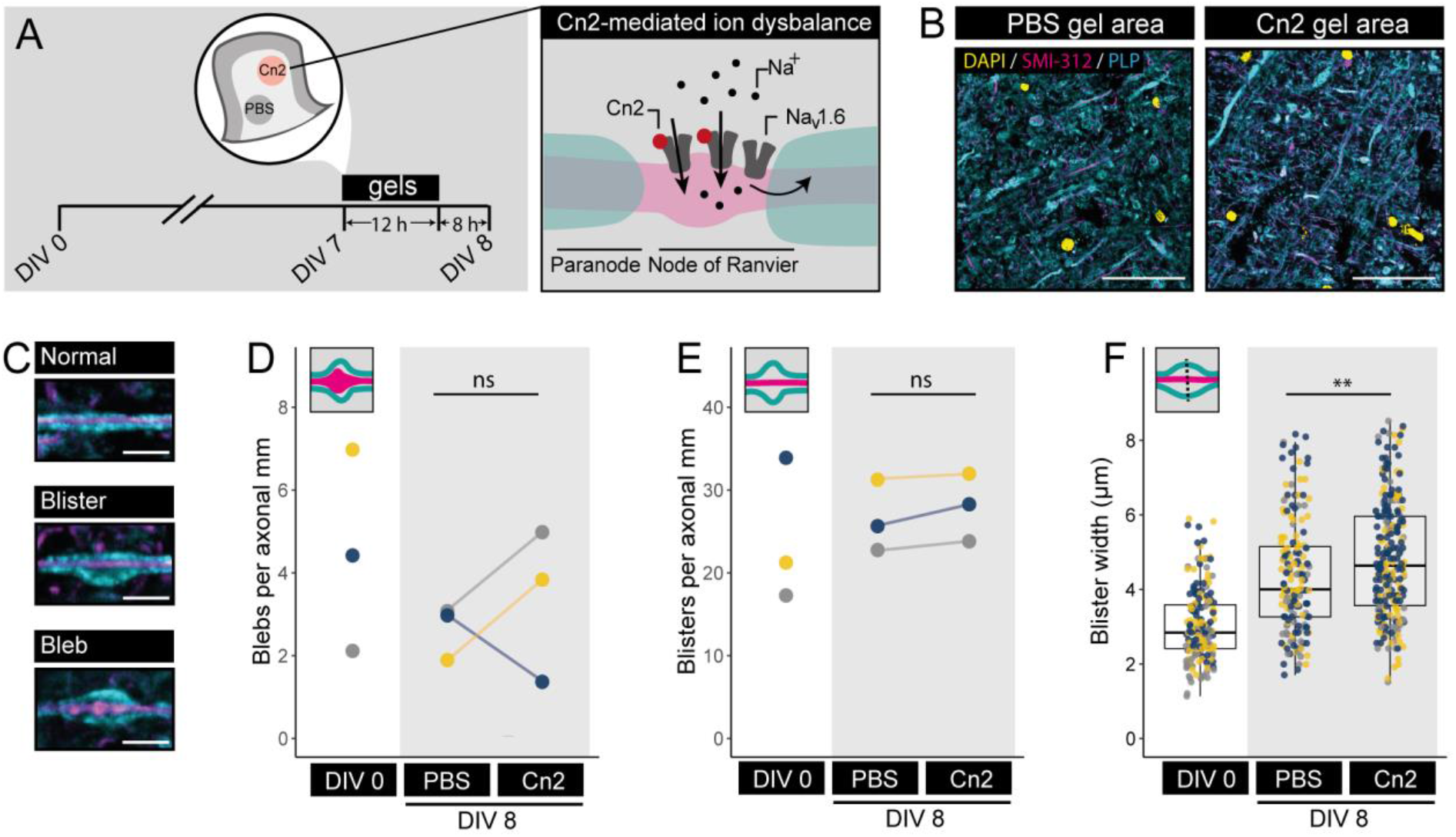
Effect of β-mammal scorpion toxin Cn2 on myelin and axon integrity. a. Schematic overview of the experimental procedure. Cryogels loaded with Cn2 were placed onto WM of HPMB-OSCs for 12 h, and myelin integrity was assessed eight hours later. b. Representative images from Cn2- and PBS-treated regions stained for DAPI (yellow), SMI-312 (magenta), and PLP (cyan). Scale bar: 50 μm. c. Representative examples of a normally myelinated axon (top), an axon with a myelin blister (middle), and an axon with axonal and myelin swelling (bottom). Scale bar: 5 μm. d. Number of axo-myelinic swellings (blebs) per axonal mm at DIV 0 and under PBS- and 140 nM Cn2-loaded gels. Biological replicates n = 3; mean analysed axonal length = 2.5 mm (range 1.6–4.4 mm). Paired t-test: DIV 0 vs PBS, t(2) = 1.06, p = 0.40; PBS vs Cn2, t(2) = 0.63, p = 0.58. e. Number of myelin blisters per axonal mm at DIV 0 and under PBS- and 140 nM Cn2-loaded gels. Biological replicates n = 3; mean analysed axonal length = 2.5 mm (range 1.6–4.4 mm). Paired t-test: DIV 0 vs PBS, t(2) = 0.44, p = 0.70; PBS vs Cn2, t(2) = 2.5, p = 0.13. f. Blister width at DIV 0 and under PBS- and 140 nM Cn2-loaded gels. Biological replicates n = 3; total blisters measured: 187 (DIV 0), 174 (PBS), 246 (Cn2). Welch’s t-test: PBS vs Cn2, t(379.7) = 2.99, p = 0.0029; mean ± SD: PBS 4.35 ± 1.51 μm, Cn2 4.81 ± 1.56 μm.

## Discussion

While organotypic slice cultures of rodent brain have proven to be a very useful tool for the modelling of pathological events in brain diseases, applying the same technology for human brains is still a challenge. Here we report on the suitability of HPMB-OSC from MS and non-MS patients for the study of myelin damage. We assessed the viability of human slices with particular emphasis on evaluating the preservation of myelin integrity throughout the culturing period. Additionally, we demonstrate the ability to induce focal and subtle myelin damage ex vivo, using cryogel-mediated delivery of LPC to produce localized demyelination and of the Na_v_1.6 agonist Cn2 to elicit myelin blister enlargement.

We found that HPMB-OSCs present gradual cellular loss with distinct temporal dynamics between GM and WM. In the GM, cellular density was decreased by DIV 9, putatively triggered by post-mortem hypoxia or axonal transection occurring during autopsy and subsequent slice preparation. In line with this, a previous study reported substantial loss of electrophysiological pyramidal neuron signatures when post-mortem delay exceeded five hours [13]. Interestingly, cell loss in the WM was not evident up to DIV13, suggesting greater resilience of WM than of GM to post-mortem insult, consistent with previous observations on subjects affected by stroke [8]. Although not quantified, cellular demise in the WM was accompanied by apparent diffuse myelin loss, as indicated by the progressive fading of the GM and WM demarcation in LFB- and PLP-stained sections. This is likely attributable to oligodendrocyte stress in response to post-mortem hypoxia and axonal transection during tissue resection. Further supporting diffuse myelin breakdown throughout the culturing period, node of Ranvier density significantly decreased by DIV 5, but stabilized after. Importantly, myelin segments that were spared did not exhibit major structural abnormalities, as indicated by the preserved nodal length up to DIV 9 and lack of paranodal intrusion into the nodal region up to DIV 13. Overall, these results indicate substantial preservation of the myelin integrity in this model.

Interestingly, spectral analysis of myelin with Nile Red solvatochromism did detect changes in WM myelin polarity throughout the culturing period, suggesting the relative stability of its molecular constituents. By contrast, GM myelin polarity significantly decreased throughout culturing. This difference may reflect regional variations in myelin composition or distinct oligodendrocyte susceptibility to hypoxic stress between GM and WM [20]. GM myelin has been suggested to primarily support the fine tuning of neural networks [36]. Consequently, GM myelin is likely to be less stable, to ensure rapid remodelling capacity during adaptive myelination.

A few important distinctions should be made from the original protocol for the acquisition of HPMB-OSC [28]. Importantly, our adaptation assumes a less strict post-mortem delay, included thicker slice cultures, and utilized antimicrobial prophylaxis after autopsy and during vibratome slicing. Although these modifications may influence tissue viability and cellular morphology, we believe these deviations have limited impacts and do not affect the generalizability of the protocol and its adaptations.

To minimize precious tissue expenditure, inter-condition variability and mimic focal pathology we utilized cryogels overlayed on individual HPMB-OSCs. Cryogels are synthetic polyethylene glycol scaffolds characterized by their relatively inertness, charge-tunability, hydrophilicity and biocompatibility [6, 25]. Previously, cryogels have been used to deliver LPC and LPS to mouse brain tissue preparations by placing them adjacent to organotypic slices [7, 40]. Here, we used cryogels which were negatively charged by partial sulphonation, which reduces their interaction with the negatively charged cell membranes, thus overall reducing risk of adherence. In our hands, delivery of fluorescently-labelled LPC onto WM via cryogels yielded focal and complete LPC penetration with limited lateral diffusion. Using this approach, we observed completely denuded axons in human tissue three days after delivery of the demyelinating agent LPC. This effect was absent when cryogels were loaded with PBS, indicating that the weight of the cryogel is unlikely to induce mechanically-evoked demyelination, like those reported by stimulation of Piezo1 receptors [37]. As such, delivery of compounds via cryogel scaffolds directly over HPMB-OSCs represents a convenient, reliable, and controllable approach in a novel human system and may provide a stepping stone for candidate drugs assessment after their initial *in vitro* evaluation. Additionally, this method opens the way to study remyelination processes ex vivo in human tissue, enabling evaluation of remyelinating strategies within a native human microenvironment.

Previously we have shown that myelin swelling and blistering occur rapidly after damage and are a hallmark of early myelin damage. Additionally, manipulation of neuronal activity was found to determine swelling fate[1, 21]. To test whether ion disbalance could induce subtle myelin damage, we focally delivered Na_v_1.6 selective stimulator β-mammal toxin Cn2 to HPMB-OSCs, which has been shown to induce persistent opening of Na_v_1.6 channels in nodal compartments [32]. While Cn2 did not increase the amount of myelin blisters, which are regarded as prodromal pathological events in MS [17, 18], Cn2 application did exacerbate the size of identified blisters in WM. The absence of more blisters after Cn2 exposure may either indicate that axon–myelin uncoupling is difficult to elicit in post-mortem cultures, or that the applied Cn2 exposure conditions (e.g. concentration of the drug and time of incubation) were insufficient to trigger additional myelin damage. Nonetheless, the increase in blister size suggests that sustained axonal sodium influx is able to drive the swelling of myelin. This is congruent with the observation that neuronal activity exacerbated myelin swelling pathology in ion-transport deficient model [19]. Interestingly, the observed increase in blister size suggests that axon-myelin coupling mechanisms remain intact in HPMB-OSCs.

Here we have investigated the suitability of HPMB-OSCs for addressing research questions pertaining to myelin. Main limitations of human-based models, such as primary cell cultures or HPMB-OSCs are the scarcity of available material and the variability between donors, influenced by factors such as medication history, comorbidities, lifestyle, or cause of death. Cryogel-mediated drug delivery circumvents these problems, as it enables focal delivery and intra-slice comparisons, thereby minimizing variability and reducing the amount of brain slices per experiment. HPMB-OSCs, in combination with cryogel-mediated drug delivery, represent a useful model to explore the mechanisms involved in the dynamics of human myelin damage and provides a powerful tool to test candidate therapies for human diseases like MS.

## Ethics approval and consent to participate

The study was approved by the local Medical Ethical Committee of the VU University Medical Center (Amsterdam, The Netherlands) and all experiments were performed in accordance with the declaration of Helsinki. For all donors, a written informed (parental) consent for a brain autopsy, the use of the medical data and permission to use the tissue for experimental purposes was obtained.

## Consent for publication

Not applicable.

## Acknowledgments

The authors are deeply grateful to the donors and their families, the autopsy team of the Netherlands Brain Bank for their services and the Mortuary staff of the Amsterdam UMC for their skillful assistance during the autopsies. We thank Dr. Tyler Kirby for providing access to and support with the spectral microscopy equipment, which was essential for this work.We thank Dr. Tyler Kirby for providing access to and support with the microscopy equipment, which was essential for this work.

## References

1. Arafa D, van de Korput J, Braaker PN, Higgins KP, Meijns NRC, Marshall-Phelps KLH, Meng J, Soong D, Scalia E, Lathem K, Keatinge M, Richmond C, Klingseisen A, Main M, Neely SA, Hampton DW, Duncan GJ, Schenk GJ, Groot ML, Chandran S, Emery B, Luchicchi A, Kole MHP, Williams AC, Lyons DA (2026) Myelin sheaths in the central nervous system can withstand damage and dynamically remodel. Science 391:eadr4661. doi: 10.1126/science.adr4661

2. Coman I, Aigrot MS, Seilhean D, Reynolds R, Girault JA, Zalc B, Lubetzki C (2006) Nodal, paranodal and juxtaparanodal axonal proteins during demyelination and remyelination in multiple sclerosis. Brain J Neurol 129:3186–3195. doi: 10.1093/brain/awl144

3. Craner MJ, Hains BC, Lo AC, Black JA, Waxman SG (2004) Co-localization of sodium channel Nav1.6 and the sodium-calcium exchanger at sites of axonal injury in the spinal cord in EAE. Brain J Neurol 127:294–303. doi: 10.1093/brain/awh032

4. Craner MJ, Lo AC, Black JA, Waxman SG (2003) Abnormal sodium channel distribution in optic nerve axons in a model of inflammatory demyelination. Brain J Neurol 126:1552–1561. doi: 10.1093/brain/awg153

5. Depp C, Sun T, Sasmita AO, Spieth L, Berghoff SA, Nazarenko T, Overhoff K, Steixner-Kumar AA, Subramanian S, Arinrad S, Ruhwedel T, Möbius W, Göbbels S, Saher G, Werner HB, Damkou A, Zampar S, Wirths O, Thalmann M, Simons M, Saito T, Saido T, Krueger-Burg D, Kawaguchi R, Willem M, Haass C, Geschwind D, Ehrenreich H, Stassart R, Nave K-A (2023) Myelin dysfunction drives amyloid-β deposition in models of Alzheimer ‘s disease. Nature 618:349–357. doi: 10.1038/s41586-023-06120-6

6. Eigel D, Schuster R, Männel MJ, Thiele J, Panasiuk MJ, Andreae LC, Varricchio C, Brancale A, Welzel PB, Huttner WB, Werner C, Newland B, Long KR (2021) Sulfonated cryogel scaffolds for focal delivery in ex-vivo brain tissue cultures. Biomaterials 271:120712. doi: 10.1016/j.biomaterials.2021.120712

7. Eigel D, Zoupi L, Sekizar S, Welzel PB, Werner C, Williams A, Newland B (2019) Cryogel scaffolds for regionally constrained delivery of lysophosphatidylcholine to central nervous system slice cultures: A model of focal demyelination for multiple sclerosis research. Acta Biomater 97:216–229. doi: 10.1016/j.actbio.2019.08.030

8. Falcao ALE, Reutens DC, Markus R, Koga M, Read SJ, Tochon-Danguy H, Sachinidis J, Howells DW, Donnan GA (2004) The resistance to ischemia of white and gray matter after stroke. Ann Neurol 56:695–701. doi: 10.1002/ana.20265

9. Gunning-Dixon FM, Brickman AM, Cheng JC, Alexopoulos GS (2009) Aging of Cerebral White Matter: A Review of MRI Findings. Int J Geriatr Psychiatry 24:109–117. doi: 10.1002/gps.2087

10. Joost S, Schweiger F, Pfeiffer F, Ertl C, Keiler J, Frank M, Kipp M (2022) Cuprizone Intoxication Results in Myelin Vacuole Formation. Front Cell Neurosci 16:709596. doi: 10.3389/fncel.2022.709596

11. van der Knaap MS, Bugiani M (2017) Leukodystrophies: a proposed classification system based on pathological changes and pathogenetic mechanisms. Acta Neuropathol (Berl) 134:351–382. doi: 10.1007/s00401-017-1739-1

12. Kocsis E, Trus BL, Steer CJ, Bisher ME, Steven AC (1991) Image averaging of flexible fibrous macromolecules: The clathrin triskelion has an elastic proximal segment. J Struct Biol 107:6–14. doi: 10.1016/1047-8477(91)90025-R

13. Kramvis I, Mansvelder HD, Meredith RM (2018) Neuronal life after death: electrophysiologic recordings from neurons in adult human brain tissue obtained through surgical resection or postmortem. Handb Clin Neurol 150:319–333. doi: 10.1016/B978-0-444-63639-3.00022-0

14. Lubetzki C, Zalc B, Williams A, Stadelmann C, Stankoff B (2020) Remyelination in multiple sclerosis: from basic science to clinical translation. Lancet Neurol 19:678–688. doi: 10.1016/S1474-4422(20)30140-X

15. Lucchinetti C, Brück W, Parisi J, Scheithauer B, Rodriguez M, Lassmann H (2000) Heterogeneity of multiple sclerosis lesions: implications for the pathogenesis of demyelination. Ann Neurol 47:707–717. doi: 10.1002/1531-8249(200006)47:6<707::aid-ana3>3.0.co;2-q

16. Luchetti S, Liere P, Pianos A, Verwer RWH, Sluiter A, Huitinga I, Schumacher M, Swaab DF, Mason MRJ (2023) Disease stage-dependent changes in brain levels and neuroprotective effects of neuroactive steroids in Parkinson ‘s disease. Neurobiol Dis 183:106169. doi: 10.1016/j.nbd.2023.106169

17. Luchicchi A, Hart B, Frigerio I, Van Dam A, Perna L, Offerhaus HL, Stys PK, Schenk GJ, Geurts JJG (2021) Axon-myelin unit blistering as early event in Normal Appearing White Matter. Ann Neurol 89:711–725. doi: 10.1002/ana.26014

18. Luchicchi A, Muñoz-Gonzalez G, Halperin ST, Strijbis E, Van Dijk LHM, Foutiadou C, Uriac F, Bouman PM, Schouten MAN, Plemel J, ‘T Hart BA, Geurts JJG, Schenk GJ (2024) Micro-diffusely abnormal white matter: An early multiple sclerosis lesion phase with intensified myelin blistering. Ann Clin Transl Neurol 11:973–988. doi: 10.1002/acn3.52015

19. Marshall-Phelps KLH, Kegel L, Baraban M, Ruhwedel T, Almeida RG, Rubio-Brotons M, Klingseisen A, Benito-Kwiecinski SK, Early JJ, Bin JM, Suminaite D, Livesey MR, Möbius W, Poole RJ, Lyons DA (2020) Neuronal activity disrupts myelinated axon integrity in the absence of NKCC1b. J Cell Biol 219:e201909022. doi: 10.1083/jcb.201909022

20. Mather ML, Evangelou AV, Bourne JN, Macklin WB, Wood TL (2025) Myelin Lipid Composition in the Central Nervous System Is Regionally Distinct and Requires Mechanistic Target of Rapamycin Signaling. Glia 73:1841–1859. doi: 10.1002/glia.70042

21. Meijns NRC, Blokker M, Idema S, Hart BA ‘t, Veta M, Ettema L, Iersel J van, Zhang Z, Schenk GJ, Groot ML, Luchicchi A (2025) Dynamic imaging of myelin pathology in physiologically preserved human brain tissue using third harmonic generation microscopy. PLOS ONE 20:e0310663. doi: 10.1371/journal.pone.0310663

22. Menichella DM, Majdan M, Awatramani R, Goodenough DA, Sirkowski E, Scherer SS, Paul DL (2006) Genetic and Physiological Evidence That Oligodendrocyte Gap Junctions Contribute to Spatial Buffering of Potassium Released during Neuronal Activity. J Neurosci 26:10984–10991. doi: 10.1523/JNEUROSCI.0304-06.2006

23. Micu I, Plemel JR, Lachance C, Proft J, Jansen AJ, Cummins K, van Minnen J, Stys PK (2016) The molecular physiology of the axo-myelinic synapse. Exp Neurol 276:41–50. doi: 10.1016/j.expneurol.2015.10.006

24. Moyon S, Frawley R, Marechal D, Huang D, Marshall-Phelps KLH, Kegel L, Bøstrand SMK, Sadowski B, Jiang Y-H, Lyons DA, Möbius W, Casaccia P (2021) TET1-mediated DNA hydroxymethylation regulates adult remyelination in mice. Nat Commun 12:3359. doi: 10.1038/s41467-021-23735-3

25. Newland B, Long KR (2022) Cryogel scaffolds: soft and easy to use tools for neural tissue culture. Neural Regen Res 17:1981–1983. doi: 10.4103/1673-5374.335156

26. Pfeiffer F, Frommer-Kaestle G, Fallier-Becker P (2019) Structural adaption of axons during de- and remyelination in the Cuprizone mouse model. Brain Pathol 29:675–692. doi: 10.1111/bpa.12748

27. Plemel JR, Michaels NJ, Weishaupt N, Caprariello AV, Keough MB, Rogers JA, Yukseloglu A, Lim J, Patel VV, Rawji KS, Jensen SK, Teo W, Heyne B, Whitehead SN, Stys PK, Yong VW (2018) Mechanisms of lysophosphatidylcholine-induced demyelination: A primary lipid disrupting myelinopathy. Glia 66:327–347. doi: 10.1002/glia.23245

28. Plug BC, Revers IM, Breur M, González GM, Timmerman JA, Meijns NRC, Hamberg D, Wagendorp J, Nutma E, Wolf NI, Luchicchi A, Mansvelder HD, Van Til NP, Van Der Knaap MS, Bugiani M (2024) Human post-mortem organotypic brain slice cultures: a tool to study pathomechanisms and test therapies. Acta Neuropathol Commun 12:83. doi: 10.1186/s40478-024-01784-1

29. Qi X, Luchetti S, Verwer RWH, Sluiter AA, Mason MRJ, Zhou J, Swaab DF (2017) Alterations in the steroid biosynthetic pathways in the human prefrontal cortex in mood disorders: A post-mortem study. Brain Pathol 28:536–547. doi: 10.1111/bpa.12548

30. Qi X-R, Verwer RWH, Bao A-M, Balesar RA, Luchetti S, Zhou J-N, Swaab DF (2019) Human Brain Slice Culture: A Useful Tool to Study Brain Disorders and Potential Therapeutic Compounds. Neurosci Bull 35:244–252. doi: 10.1007/s12264-018-0328-1

31. Ransohoff RM (2012) Animal models of multiple sclerosis: the good, the bad and the bottom line. Nat Neurosci 15:1074–1077. doi: 10.1038/nn.3168

32. Schiavon E, Sacco T, Cassulini RR, Gurrola G, Tempia F, Possani LD, Wanke E (2006) Resurgent current and voltage sensor trapping enhanced activation by a beta-scorpion toxin solely in Nav1.6 channel. Significance in mice Purkinje neurons. J Biol Chem 281:20326–20337. doi: 10.1074/jbc.M600565200

33. Stadelmann C, Timmler S, Barrantes-Freer A, Simons M (2019) Myelin in the Central Nervous System: Structure, Function, and Pathology. Physiol Rev 99:1381–1431. doi: 10.1152/physrev.00031.2018

34. Teo W, Caprariello AV, Morgan ML, Luchicchi A, Schenk GJ, Joseph JT, Geurts JJG, Stys PK (2021) Nile Red fluorescence spectroscopy reports early physicochemical changes in myelin with high sensitivity. Proc Natl Acad Sci 118:e2016897118. doi: 10.1073/pnas.2016897118

35. Teo W, Morgan ML, Stys PK (2025) Quantitation of the physicochemical properties of myelin using Nile Red fluorescence spectroscopy. J Neurochem 169:e16203. doi: 10.1111/jnc.16203

36. Timmler S, Simons M (2019) Grey matter myelination. Glia 67:2063–2070. doi: 10.1002/glia.23614

37. Velasco-Estevez M, Gadalla KKE, Liñan-Barba N, Cobb S, Dev KK, Sheridan GK (2020) Inhibition of Piezo1 attenuates demyelination in the central nervous system. Glia 68:356–375. doi: 10.1002/glia.23722

38. Verwer RWH, Baker RE, Boiten EFM, Dubelaar EJG, van Ginkel CJM, Sluiter AA, Swaab DF (2003) Post-mortem brain tissue cultures from elderly control subjects and patients with a neurodegenerative disease. Exp Gerontol 38:167–172. doi: 10.1016/s0531-5565(02)00154-7

39. Verwer RWH, Hermens WTJMC, Dijkhuizen P, ter Brake O, Baker RE, Salehi A, Sluiter AA, Kok MJM, Muller LJ, Verhaagen J, Swaab DF (2002) Cells in human postmortem brain tissue slices remain alive for several weeks in culture. FASEB J Off Publ Fed Am Soc Exp Biol 16:54–60. doi: 10.1096/fj.01-0504com

40. Walsh CM, Hill S, Newland B, Dooley D (2025) Cryogel scaffolds for localised delivery of lipopolysaccharide in organotypic spinal cord slice cultures: A novel ex vivo model of neuroinflammation. Mater Today Bio 34:102211. doi: 10.1016/j.mtbio.2025.102211

41. Zoupi L, Booker SA, Eigel D, Werner C, Kind PC, Spires-Jones TL, Newland B, Williams AC (2021) Selective vulnerability of inhibitory networks in multiple sclerosis. Acta Neuropathol (Berl) 141:415–429. doi: 10.1007/s00401-020-02258-z

42. Zuroff L, Farkhondeh V, Bove R, Green AJ (2025) The Road to Remyelination in Multiple Sclerosis: Breakthroughs, Challenges, and Considerations for Future Trial Design. Drugs. doi: 10.1007/s40265-025-02212-x

